# Convergent regulatory evolution and the origin of flightlessness in palaeognathous birds

**DOI:** 10.1101/262584

**Authors:** Timothy B. Sackton, Phil Grayson, Alison Cloutier, Zhirui Hu, Jun S. Liu, Nicole E. Wheeler, Paul P. Gardner, Julia A. Clarke, Allan J. Baker, Michele Clamp, Scott V. Edwards

**Author notes:** correspondence to: TBS or SVE.

## Abstract

The relative roles of regulatory and protein evolution in the origin and loss of convergent phenotypic traits is a core question in evolutionary biology. Here we combine phylogenomic, epigenomic and developmental data to show that convergent evolution of regulatory regions, but not protein-coding genes, is associated with flightlessness in palaeognathous birds, a classic example of a convergent phenotype. Eleven new genomes, including a draft genome from an extinct moa, resolve palaeognath phylogeny and show that the incidence of independent, convergent accelerations among 284,000 conserved non-exonic elements is significantly more frequent in ratites than other bird lineages. Ratite-specific acceleration of conserved regions and measures of open chromatin across eight tissues in the developing chick identify candidate regulatory regions that may have modified or lost function in ratites. Enhancer activity assays conducted in the early developing chicken forelimb confirm that volant versions of a conserved element in the first intron of the *TEAD1* gene display conserved enhancer activity, whereas an accelerated flightless version fails to drive reporter gene expression. Our results show that convergent molecular changes associated with loss of flight are largely regulatory in nature.

---

Convergent evolution – the independent evolution of similar phenotypes in divergent taxa – produces some of the most striking examples of adaptation, but the molecular architecture of convergent traits is not well understood (Losos 2011; Martin and Orgogozo 2013; Stern 2013; Storz 2016). In some cases, similar or identical mutations in independent lineages appear to be associated with convergent phenotypes (Chan et al. 2010; Zhen et al. 2012), whereas in other cases convergent phenotypes appear to arise by diverse molecular paths (Cooper et al. 2014). While studies of convergence are increasingly focused on detecting genome-wide patterns (Foote et al. 2015; Marcovitz et al. 2016; Yeaman et al. 2016; Partha et al. 2017; Xu et al. 2017), prior work often has been biased towards protein-coding regions, and it remains an open question whether regulatory regions are more or less likely to be involved in convergent phenotypes than protein-coding genes.

A particularly notable example of a convergent trait involves loss of powered flight, which has occurred many times independently in the course of avian evolution. One of the most iconic groups of flightless birds is the ratites, consisting of extant ostrich, kiwis, rheas, cassowaries, and emu, and the extinct moas and elephant birds. All ratites show morphological similarities including forelimb reduction (ranging from moderate in ostrich and rheas to a complete absence in moas), reduced pectoral muscle mass associated with the absence of the sternal keel, and feather modifications, as well as generally larger body size (Houde 1986; Bickley and Logan 2014; Nesbitt and Clarke 2016). Historically, ratites were thought to be a monophyletic sister clade to the volant tinamous (Cracraft 1974), but, despite remaining uncertainties in palaeognath relationships, recent molecular phylogenetic evidence strongly supports ratite paraphyly, implying as many as six independent losses of flight within this group based on these data and biogeographic scenarios (Harshman et al. 2008; Baker et al. 2014; Mitchell et al. 2014; Nesbitt and Clarke 2016; Grealy et al. 2017; Yonezawa et al. 2017). Recent developmental work in ratites has focused on protein-coding regions underlying limb reduction (de Bakker et al. 2013; Bickley and Logan 2014; Farlie et al. 2017), and a recent study of loss of flight in Galapagos cormorants (Burga et al. 2017), while identifying putative genic drivers of flightlessness, did not focus on possible roles of the noncoding genome in loss of flight (although see (Berger and Bejerano 2017)). The relative roles of protein-coding versus regulatory change (King and Wilson 1975) in loss of flight, in ratites or other birds, is unknown.

To study the genomic correlates of flight loss in ratites, we assembled and annotated 11 new palaeognath genomes (Methods Section 1, Supplemental Table 1, Supplemental Table 2), including 8 flightless ratites (greater rhea, lesser rhea, cassowary, emu, Okarito kiwi, great spotted kiwi, little spotted kiwi, and the extinct little bush moa (Baker et al. 2014; Cloutier et al. 2018b)) and 3 tinamous (thicket tinamou, elegant crested tinamou, and Chilean tinamou), and analyzed them together with the published ostrich (Jarvis et al. 2014), white-throated tinamou (Jarvis et al. 2014), and North Island brown kiwi (Le Duc et al. 2015) genomes. We generated new transcriptome data from three tissues (brain, liver, gonad) from both emu and Chilean tinamou, annotated between 17,021 and 21,342 gene models (Methods Sections 2 and 3, Supplemental Figure 1, Supplemental Table 3) in these genomes and identified 284,001 conserved non-exonic elements (CNEEs) based on a novel, phylogenetically informed (Paten et al. 2011) whole genome alignment including 35 birds and 7 non-avian reptiles (Methods Section 4), facilitating evolutionary and functional analysis of the molecular basis of convergence in ratites.

## Incomplete lineage sorting explains persistent challenges of palaeognath phylogenetics

We compiled a total evidence data set of 41,184,181 bp of aligned DNA from 20,850 noncoding loci, including introns, ultraconserved elements (UCEs) (McCormack et al. 2012) and CNEEs (Edwards et al. 2017) (Supplemental Table 4, Methods Section 5) to test the hypothesis of ratite paraphyly and to resolve the placement of rheas, which remains an outstanding question in palaeognath phylogenetics (Harshman et al. 2008; Phillips et al. 2010; Yonezawa et al. 2017).

Consistent with recent molecular phylogenies (Harshman et al. 2008; Phillips et al. 2010; Baker et al. 2014; Mitchell et al. 2014; Yonezawa et al. 2017), we recover a basal divergence between the ostrich and the remaining palaeognaths, including the tinamous, which are therefore nested within a paraphyletic ratite clade (Figure 1A). However, contrary to recent concatenation analyses (Yonezawa et al. 2017), our coalescent (Liu et al. 2010) analysis consistently places the rheas as sister to the kiwi+emu+cassowary clade. Our species tree is robust to a wide range of alignment, tree-building and data filtering choices (Supplemental Figure 2), is recovered stably with maximum support in coalescent analyses (Figure 1A, Supplemental Figure 3), and is corroborated by 4274 informative CR1 retroelement insertions, including 18 absent in ostrich but shared among all non-ostrich palaeognaths, including tinamous (Cloutier et al. 2018a). In contrast, no retroelement insertions support ratite monophyly, and the sequence data strongly rejects ratite monophyly using multiple methods (Cloutier et al. 2018a).

**Figure 1.**
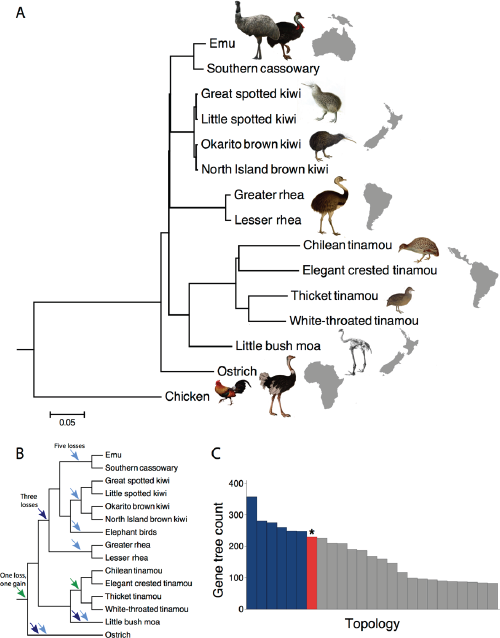
Genome-wide data places rheas as sister to a kiwi/cassowary-emu clade. A) MP-EST species tree topology, with 100% bootstrap support throughout (not shown). Branch lengths in units of substitutions per site were estimated with ExaML using a fully partitioned alignment of 20,850 loci constrained to the MP-EST topology. B) Ratite paraphyly is consistent with either a single loss of flight in the palaeognath ancestor followed by regain in tinamous (green arrows) or a minimum of three independent losses of flight in the ratites (dark blue arrows), with six losses suggested by a proposed sister group relationship between elephant birds and kiwi (Mitchell et al. 2014) (light blue arrows). C) Distribution of the 25 most common gene tree topologies, showing that the species tree topology (in red and marked with a star symbol) is not the most common gene tree topology.

Our analysis and taxon sampling implies a minimum of three losses of flight in ratites, while the inferred biogeographic history considering all extant and extinct ratites has been used to argue for up to six losses of flight in the history of this group (Figure 1B), with independent losses of flight in the ancestors of rhea, kiwi, and the emu+cassowary clade (Mitchell et al. 2014; Grealy et al. 2017; Yonezawa et al. 2017). The alternative – a single loss of flight at the base of the palaeognaths followed by a regain of flight in tinamous – appears implausible given evidence for repeated losses of flight across birds and the lack of any evidence for regains of flight after loss.

The species tree is characterized by successive short internal branches leading to the common ancestor of kiwi and emu+cassowary, and the common ancestor of this clade with rheas (Figure 1A, Supplemental Figure 4). Incomplete lineage sorting (ILS) across short internodes can produce topologies for individual genes that differ from the species tree, and in the extreme case can result in an anomaly zone where the most common gene tree discords with the species tree (Degnan and Rosenberg 2006). Empirical anomaly zones are thought to be rare (Linkem et al. 2014), but we find evidence for such a zone in palaeognaths, wherein the gene tree that matches the species tree is less common than other gene trees (Figure 1C). Median bootstrap support for conflicting clades in individual gene trees is moderate, but there is strong relative likelihood support for alternative gene tree topologies in the data set (Supplemental Figure 5), suggesting that these results do not arise solely from uninformative gene trees. Estimated coalescent branch lengths are also consistent with values expected to produce anomalous gene trees across the observed short internal branches (Degnan and Rosenberg 2006; Rosenberg 2013) (Supplemental Figure 4). Together, these results suggest that palaeognaths fall within an empirical anomaly zone and that substantial ILS has contributed to past difficulties in resolving relationships within this group (Mitchell et al. 2014; Yonezawa et al. 2017; Cloutier et al. 2018a).

## Little evidence for convergent evolution of protein-coding genes

Prior work on the molecular basis of convergent phenotypes has focused on amino acid substitutions that occur independently on multiple lineages that share a convergent phenotype (Zhen et al. 2012; Natarajan et al. 2016; Xu et al. 2017), a process that is also a common outcome of neutral processes (Castoe et al. 2009; Zou and Zhang 2015) and likely not a major determinant of patterns of evolution genome-wide (Thomas and Hahn 2015). However, to test for an excess of convergent amino acid substitutions between ratite lineages, we analyzed 6,337 high quality single-copy orthologs present in at least 33 of 35 bird species in our alignment (including all available palaeognaths except the little bush moa and North Island brown kiwi (Le Duc et al. 2015) due to lower quality protein annotations in those species; Methods Section 6). The number of convergent substitutions in ratite-ratite branch pairs can be predicted solely from neutral branch lengths (Zou and Zhang 2015) and the number of divergent substitutions, with no evidence for a significant effect of a ratite-specific model term (Supplemental Table 5). These results are robust to a variety of analysis choices, including using gene trees instead of species trees (Mendes and Hahn 2016), strict data filtering, and non-linear parameterizations (Supplemental Table 5).

An arguably more likely form of protein evolution underlying convergent phenotypes is consistent rate shifts due to lineage-specific positive selection or changes in constraint in phenotypically convergent lineages (Chikina et al. 2016; Partha et al. 2017). To detect such shifts, we computed normalized amino acid branch lengths for 6,436 single-copy orthologs (Chikina et al. 2016), and generated empirical P-values (for acceleration and deceleration separately) based on 1,994 randomly permuted datasets (Methods Section 6). After FDR correction, we find no cases of proteins exhibiting convergently increased or decreased rates of evolution in ratite lineages, suggesting little evidence for a common role of such protein shifts in the evolution of flightless phenotypes.

This result does not arise because we lack power to detect convergent rate shifts: consistent with earlier work (Zhang et al. 2014), we find 150 convergently accelerated and 332 convergently decelerated genes (at a 5% FDR, using empirical P-values) in convergent lineages capable of vocal learning (hummingbirds, parrots, and songbirds, representing either 2 or 3 gains). However, we find no evidence for any GO enrichments among either the accelerated or decelerated gene sets, and we likely overestimate the number of convergent rate shifts due to the overrepresentation of fast-evolving passerines relative to other vocal learners in our vocal learning species set and our tendency to identify passerine-specific patterns as more general ‘vocal learning’ patterns. Nonetheless, we find that vocal learners, but not ratites, have an unusually high number of inferred convergent rate shifts compared to random trios of birds, even after excluding trios of species with more than one passerine (Figure 2A; permutation P-value (accelerated) = 0.02, permutation P-value (decelerated) = 0.028). We also identify highly correlated estimates of clade-specific rate shifts using an alternate method, the RELAX test in HyPhy (Wertheim et al. 2015), which suggests these results are robust to analysis methods (Supplemental Figure 6).

**Figure 2.**
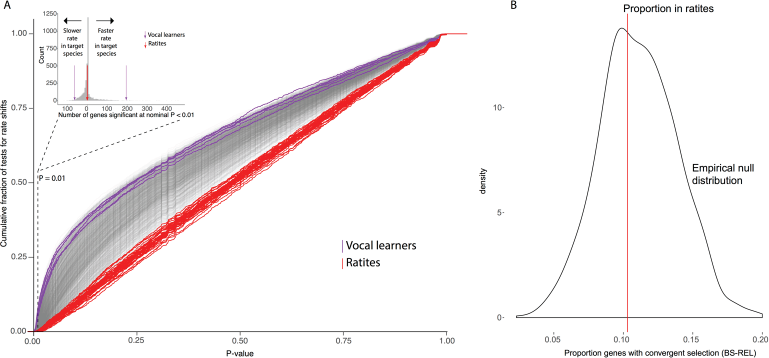
Minimal evidence for protein-coding convergence in ratites. A) Empirical cumulative P-value distributions for the test for convergent rate shifts in specific avian lineages for random trios (gray), ratite trios (red), and vocal learner trios (purple). Inset shows a histogram of the count of nominally significant rate shifts at P < 0.01 for random trios of non-ratite, non-vocal learning, non-sister birds. Observed mean is show for ratite trios (red arrows) and vocal learning trios (purple arrows). B) Empirical null distribution of expected proportion of lineages with evidence for convergent positive selection (defined as selection in at least 2 lineages and 0 non-target lineages), based on 1000 simulations. The observed value in ratites (red line) is not significantly different from the null expectation (permutation P = 0.6).

Positive selection on a subset of codons within a gene may not be detectable using relative rate tests, but still could be of major functional importance to phenotypic evolution and convergence. We find 301 genes that are uniquely selected in ratites, but the number (31, or 10.3%) that are selected in at least two ratite species not greater than expected by chance (permutation P = 0.60; Figure 2B), and neither the set exclusive to ratites nor this convergent subset are enriched for any GO terms (P > 0.05 for all categories after multiple test correction). Nonetheless, we do find two genes with evidence for positive selection in at least three ratites, but no other species: *ZFHX4* and *MGST2*. While neither gene is extensively characterized in chickens, *ZFHX4* is a transcription factor expressed in the forelimb AER during chicken development (Darnell et al. 2007), while *MGST2* is expressed in flight feathers (Ng et al. 2015).

Another potential molecular consequence of phenotypic convergence is repeated loss or pseudogenization of identifiable protein-coding genes across one or more ratite lineages (Hiller et al. 2012; Meredith et al. 2014). To conservatively test for this scenario, we used blastp to screen all chicken proteins (Ensemble release 86) against predicted proteins in each palaeognath species as well as three outgroup non-avian reptiles (green anole, American alligator, and painted turtle) and 27 neognaths. We find no cases of a gene lost in all ratites, although this measure will generally fail to detect partially degenerated pseudogenes.

As a less conservative measure for divergence and possible modification of protein function, we applied a profile hidden Markov model based method to detect sequence changes that are underrepresented in homologs of a protein (Wheeler et al. 2016), an approach similar to that used to identify putative function-altering mutations in protein-coding genes associated with loss of flight in the Galapagos cormorant (Burga et al. 2017). Only a single gene, *Neurofibromin 1* (*NF1*), shows evidence for convergent functional evolution in ratites by this measure (at a 5% FDR), while 40 genes show evidence for convergent functional evolution in vocal learning species (at a 5% FDR). In neither ratites nor vocal learners do we detect an excess of convergent function-alternating protein-coding substitutions compared to genome-wide distributions (Supplemental Figure 7). Little is known about *NF1* function in chickens, but in mice it has been shown to be involved in skeletal (Kolanczyk et al. 2007) and skeletal muscle development (Kossler et al. 2011).

Overall, although we detect a small number of possible candidate genes exhibiting patterns of protein evolution consistent with a role in convergent phenotypic evolution, the genome-wide impacts of these signals appear small, and always less than observed in vocal learners or random bird lineages. Taken together, these results suggest that proteins are unlikely to be major drivers of convergent phenotypes across the multiple independent losses of flight in ratites.

## Conserved non-exonic elements are convergently accelerated in ratites

Regulatory regions may be subject to less pleiotropic constraint than protein-coding genes (Carroll et al. 2008; Chan et al. 2010), and thus convergent regulatory evolution may be more likely than that in protein-coding genes if common pathways are involved. To identify candidate regulatory regions in palaeognath genomes, we identified 284,001 CNEEs using PHAST (Siepel et al. 2005), which are thought to have regulatory roles in birds (Seki et al. 2017) and other taxa (Pennacchio et al. 2006; Visel et al. 2008; Capra et al. 2013). We used a novel Bayesian method (Hu et al. 2018) to model changes in conservation of these elements across the phylogeny.

We identified 2,053 ratite-accelerated regions (RARs), defined as CNEEs with strong evidence for acceleration (loss of constraint) in one or more ratite lineage and constraint in all other birds (Figure 3A). Of these, 1,683 (81.2%) are also identified as ratite-accelerated (but not tinamou-accelerated) by phyloP (Pollard et al. 2010). These elements represent strong candidates for regions of the genome with altered function in ratite lineages, although we do not attempt to distinguish between mutational processes (e.g., GC-biased gene conversion), positive selection for function-alternating substitutions, or neutral accumulation of mutations subsequent to loss of constraint (but see (Hu et al. 2018)). To detect convergent changes in constraint, we used the posterior probabilities of conservation and acceleration at each node to estimate the posterior expected number of independent accelerations for each CNEE. Among the 2,053 RARs, up to 839 (40.8%) have experienced two or more independent acceleration events, dropping to 399-586 with additional filtering (Supplemental Figure 8).

**Figure 3.**
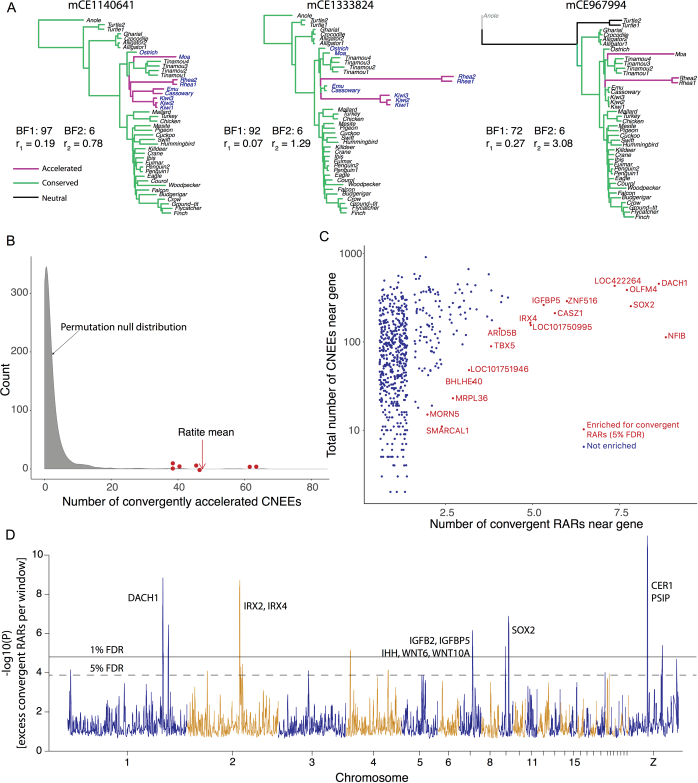
Ratites exhibit unusually high numbers of convergently accelerated CNEEs. A) Trees for three examples of convergent RARs. Branch lengths are proportional to substitution rate relative to neutrality and colored according to the maximum *a posteriori* conservation state. Bayes Factor 1 and 2 (see main text), as well as the conserved (r1) and accelerated (r2) rates are indicated for each element. B) Density plot of the null distribution of the observed number of convergently accelerated elements in random trios of neognaths (gray), with individual ratite trios (red points) and ratite mean (red arrow) indicated; ratites have significant excess of convergence (permutation P-value = 0.0063). C) Genes with evidence for excess convergent RARs. Each point represents the number of CNEEs associated with a gene and the number of convergent RARs associated with that gene. Colored and labeled points are genes with excess convergent RARs based on a permutation test (5% FDR). Only genes with at least one convergent RAR are plotted. D) Evidence for spatial clustering of convergent RARs across the genome. The y-axis shows the negative log10 p-value for a test of excess convergent RARs in 1 Mb sliding windows (100kb slide), computed using a binomial test, where the probability of success is the total RARs / total CNEEs, and the number of samples is the number of CNEEs in each window. Genes in each window that may be of particular interest are noted.

To determine if ratites experience more convergent acceleration in CNEEs than is typical for birds, we randomly sampled trios of non-sister neognaths and counted how many CNEEs were convergently accelerated in all three lineages using the posterior probabilities from the unrestricted Bayesian model. Among random trios of neognaths, we observe a mean of 2.7 convergently accelerated CNEEs (across 1,419 permutations). In contrast, we observe a mean of 47.3 convergently accelerated CNEEs among seven ratite trios (conservatively requiring acceleration in both moa and ostrich plus at least one additional clade). The mean number of convergently accelerated CNEEs we observe in ratites is thus unusually high compared to an empirical null distribution (permutation P-value = 0.0063, Figure 3B).

We next tested whether RARs and/or convergent RARs are associated with particular functional annotations, genes, or chromosomal locations. Using a permutation test that accounts for the non-random distribution of CNEEs across the genome, we find evidence that convergent RARs are enriched for an association with transcription factor ontology terms (Table 1). Moreover, we identify 17 genes (including 8 transcription factors: DACH1, IRX4, TBX5, SOX2, NFIB, BHLHE40, CASZ1, ZNF516) associated with a larger than expected number of convergent RARs (at a 5% FDR, by permutation; Figure 3C), and 21 genes (including 10 transcription factors and co-factors: DACH1, TBX5, VSX1, SOX2, NFIB, CASZ1, NPAS3, ARID5B, HOXD4, BAZ1A) associated with a larger than expected number of RARs (regardless of convergence; Supplemental Figure 9). Notably, TBX5, an essential transcription factor for both forelimb and heart development, has been previously associated with the delay in embryonic forelimb outgrowth displayed by emu; first through a proposed delay in the onset of TBX5 expression in the lateral plate mesoderm (Bickley and Logan 2014) and more recently as a potential regulator for NKX2.5 (Farlie et al. 2017). The large number of convergent RARs we observe associated with TBX5 suggests that its regulatory landscape may be a common target of selection or relaxation of constraint during transitions to avian flightlessness.

**Table 1:** Gene ontology terms enriched in genes near convergent RARs.

| Ontology | Term(*) | Target<br>Fraction | Exp. Target<br>Fraction | P-value | Q-value |
| --- | --- | --- | --- | --- | --- |
| MF | DNA binding | 0.4408602 | 0.3198529 | 0.008357771 | 0.0292522 |
| MF | nucleic acid binding | 0.4946237 | 0.3629177 | 0.006060606 | 0.0292522 |
| BP | regulation of gene expression | 0.4705882 | 0.3999947 | 0.04582909 | 0.1920457 |
| BP | regulation of nucleic acid-templated transcription | 0.4411765 | 0.3660169 | 0.03575788 | 0.1830722 |

\*selected non-redundant terms from each ontology; full list in supplemental data

Computational approaches for assigning CNEEs to the genes they regulate are necessarily imprecise, so we also looked for genomic regions (irrespective of gene annotations) that are enriched for RARs or convergent RARs. We find 11 regions of the genome with an excess of convergent RARs (at a 5% FDR; Figure 3D), and an additional 13 windows with an excess of RARs (irrespective of convergent signal). These include regions centered on known regulators of limb development (e.g., DACH1) and on regions near genes associated with body size (IGFBP2, IGFBP5). This latter region corresponds to a gene desert previously shown to be enriched in avian-specific CNEEs (Lowe et al. 2015). Overall, these analyses demonstrate that convergently accelerated CNEEs in ratites are enriched near transcription factors and other genes that may be associated with convergent phenotypes.

## Functional characterization of ratite-accelerated CNEEs

RARs are candidates for regulatory regions associated with convergent phenotypes in ratites, including loss of flight. To functionally characterize the association between sequence conservation and regulatory activity in birds, we used ATAC-seq (Buenrostro et al. 2013) to identify genomic regions of open chromatin for 8 tissues during the course of chick development. ATAC-seq peaks have been demonstrated to be enriched for active regulatory regions and highly enriched for transcription start sites (TSS) across diverse tissues and species (Buenrostro et al. 2013; Gehrke et al. 2015). As expected, the ATAC-seq landscape was strongly enriched for chicken transcription start sites for all eight tissues (min = 7.95x, max = 11.09x, mean = 9.70x, p < 0.001 for all samples, Supplemental Figure 10A), and also for ChIP-seq peaks identified in a previous chicken developmental study (Seki et al. 2017) (Supplemental Figure 10B). Consistent with recent work (Seki et al. 2017), CNEEs are also enriched under ATAC-seq peaks (min = 1.35x, max = 2.15x, mean = 1.77x, p < 0.001 for all samples, Figure 4A), further supporting the regulatory function of CNEEs in this and other datasets.

**Figure 4.**
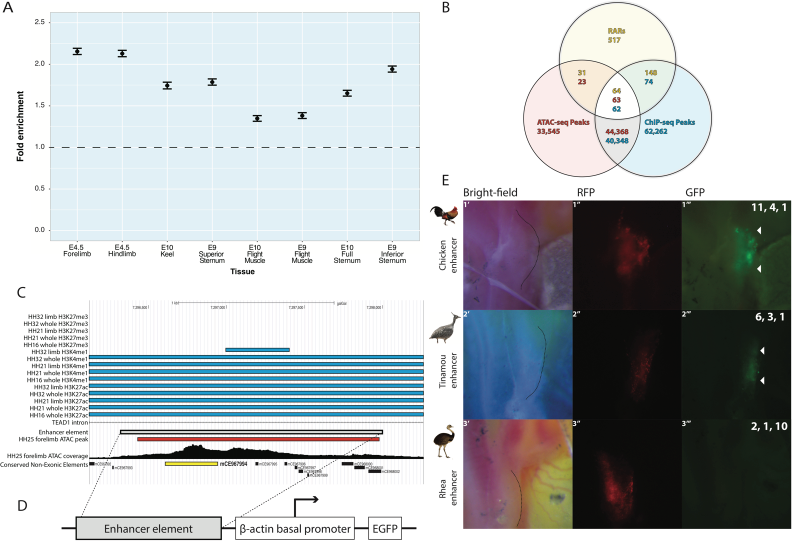
Ratite-specific convergent acceleration is associated with altered activity in a candidate enhancer. A) CNEEs are enriched under ATAC-seq peaks across all eight chicken tissues. Enrichment fold change (mean and 95% CI), calculated against chicken genomic background using GAT (10,000 samples). B) Candidate forelimb ATAC-seq peaks for enhancer screen were identified by overlap to convergent RARs (that were additionally supported by phyloP tests) and ChIP-seq peaks (Seki et al. 2017). C) Genomic region associated with strong enhancer activity in early chicken forelimbs. D) Schematic of B-actin/GFP vector used in enhancer screen. E) Results of enhancer screen of candidate ATAC-seq peak in chicken forelimb bud. Pictures are representative images of 10-16 replicates showing developing forelimb region 24 hours after electroporation (HH19-20). Dashed line on the bright field image (left column) indicates the forelimb region. RFP expression (red, middle column) indicates the area electroporated. GFP expression (green, right column) is driven by enhancer activity of the candidate region. Number of replicates showing strong, partial, and no GFP expression are indicated in upper right corner of each image. e1) Chicken ATAC-seq peak drives consistent GFP expression throughout the developing forelimb (strong GFP 11/16, partial GFP 4/16, no GFP 1/16). e2) The homologous genomic region in the elegant crested tinamou also drives consistent GFP expression throughout the developing chicken forelimb (strong GFP 6/10, partial GFP 3/10, no GFP 1/10). e3) The homologous genomic region in the greater rhea fails to drive consistent GFP expression throughout the developing chicken forelimb (strong GFP 2/13, partial GFP 1/13, no GFP 10/13).

To test whether ratite-specific sequence acceleration results in functional differentiation of enhancer activity *in vivo*, we screened a set of candidates for enhancer activity in developing chicken forelimbs. We focused on putative forelimb enhancers identified by examining the intersection of consistent forelimb ATAC-seq peaks, convergent RARs, and previously published embryonic chicken ChIP-seq peaks (Seki et al. 2017), resulting in 63 candidate enhancers (Figure 4B). Using an electroporated beta-actin/GFP enhancer construct assay, we identified a promising chicken region from among these candidates, consisting of the ATAC-seq peak associated with convergent RAR mCE967994 which produced consistent strong enhancer activity in early chicken forelimbs (Figure 4CE). We tested for enhancer activity in the homologous genomic region in the volant elegant crested tinamou, in which mCE967994 is identified as conserved in our Bayesian analysis, and the flightless greater rhea, in which mCE967994 is accelerated. We find that the tinamou version of this enhancer consistently drives GFP expression (N = 9 of 10 embryos), but the rhea version does not (Figure 4E) (N = 10 of 13 embryos). Thus, accelerated sequence evolution of this element in rheas appears associated with functional divergence of regulatory activity.

## Conclusions

Understanding the relative roles of regulatory and protein-coding change in phenotypic evolution (King and Wilson 1975), as well as the role of convergent genomic mechanisms in the evolution of convergent phenotypes, are longstanding questions in evolutionary biology. Here, unbiased statistical and functional screens indicate convergent evolutionary changes associated with flightlessness in ratites are primarily regulatory in nature. While we do not rule out a role for protein-coding changes in morphological evolution in these species, we suggest that protein-coding evolution, in contrast to regulatory evolution, must be largely lineage-specific. Our results contrast with previous work examining the genomic correlates of flightlessness in birds, which has focused solely on protein-coding regions (Burga et al. 2017) and transcriptome phenotypes, in some cases implying regulatory evolution without directly testing it (de Bakker et al. 2013; Bickley and Logan 2014; Farlie et al. 2017). Future work should increase attention to the possible role of noncoding regulatory regions in the origin of flightlessness in birds.

## Data availability

All sequencing data generated in this project is available at NCBI under BioProjects PRJNA433110 (genomes), PRJNA433154 (ATAC-seq), and PRJNA433114 (RNA-seq). In addition, processed data is available from Dryad; an alignment hub for the UCSC genome browser, and affiliated files, can be accessed at <URL TO BE DETERMINED>.

## Code availability

All scripts used for data processing and analysis are available on Github at https://github.com/tsackton/ratite-genomics. Further details are provided in the supplemental methods.

## Acknowledgements

We gratefully acknowledge the Ngati Hine, Ngati Hei, Tuhoe, Ngati Tuwharetoa, Ngati Raukawa, and Ngai Tahu peoples, who permitted genetic analyses of kiwi blood samples obtained from their lands. We thank Carol Ritland, Myles Lamont, Kevin Kerr, Graham Crawshaw, Dan Janes, Rob Fleischer, and Jeremiah Trimble for assistance with sample acquisition. Cliff Tabin, John Young, and members of the Tabin and Cepko labs provided advice, assistance, training, and equipment that enabled the developmental work reported here. Benedict Paten and Joel Armstrong provided assistance with progressiveCactus. Aaron Kitzmiller and James Cuff provided advice and support on computational and software issues. We also thank Terry Capellini and Nathan Clark for comments and discussion. The computations in this paper were run on the Odyssey cluster supported by the FAS Division of Science, Research Computing Group at Harvard University, the Sanger Institute Computing Cluster, and the Abacus Computing Cluster at Canterbury University. This project was supported by NSF grant 1355343 to JAC, AJB, MC, and SVE. PG was supported by an NSERC PGSD-3 grant.

## Author contributions

JAC, AJB, and SVE conceived this project with help from MC. SVE provided overall project leadership. AJB, AC, and SVE obtained samples. PG and AC prepared DNA, RNA, and sequencing libraries. TBS, SVE, and MC designed analysis strategies with assistance from all authors. TBS provided supervision for all bioinformatics, assembled and annotated all genomes, made the whole genome alignment, identified conserved elements, identified homologous proteins, and led the analysis of protein-coding evolution, non-coding evolution, and convergence. AC compiled protein, CNEE, UCE, and intron alignments and led the phylogenetics analysis. PG prepared samples for ATAC-seq, analyzed ATAC-seq data with assistance from TBS, and carried out all developmental work. NEW and PPG developed and applied the DeltaBS method to detect functional divergence. ZH and JSL developed and applied the phyloAcc method. TBS wrote the manuscript together with AC, PG, and SVE. All authors edited the manuscript and approved the final draft.

**Supplemental Figures and Tables follow references cited.**

---

**Supplemental Figure 1.**
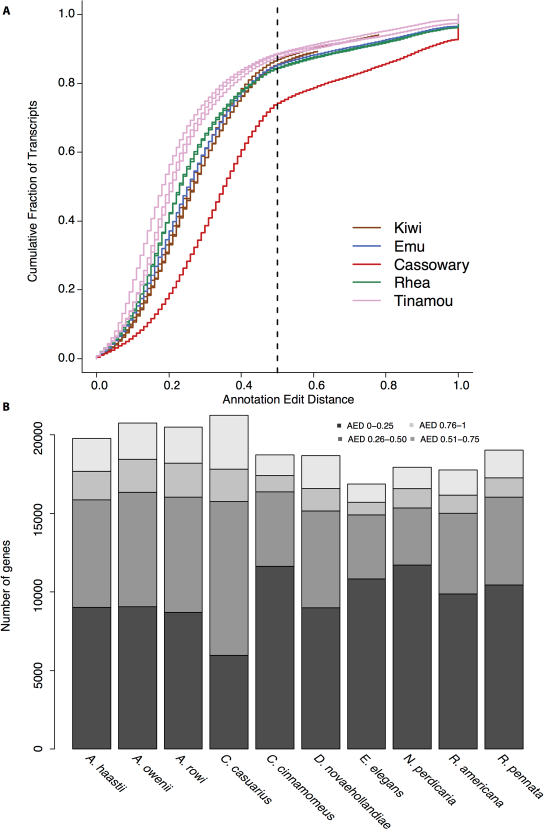
Genome annotations are of high quality in all newly sequenced species. A) Distribution of annotation edit distances from MAKER annotations across our genomes, which generally show acceptable to high quality results. B) Number of genes annotated across each species, in bins based on annotation edit distance. Lower numbers indicate better agreement with evidence (RNA-seq and cross species protein mapping). In all cases, a substantial majority of genes are annotated with AED below 0.5.

**Supplemental Figure 2.**
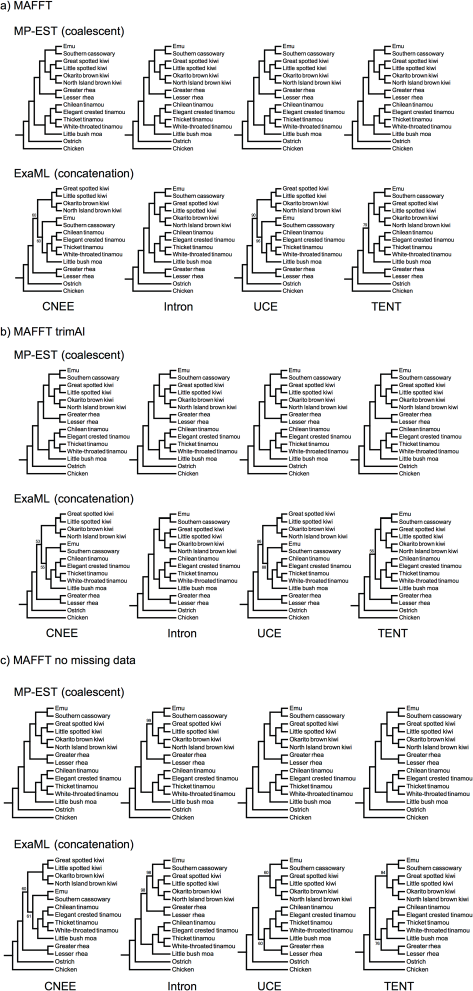
Inference of palaeognath relationships from MP-EST coalescent based and ExaML concatenation analyses. Topologies are given for separate analyses of conserved non-exonic elements (CNEEs), introns, and ultraconserved elements (UCEs), as well as for the total evidence nucleotide tree (TENT) combining loci from all three marker types. Only bootstrap support values < 100% are drawn. Topologies are shown for MAFFT sequence alignments with no additional trimming or filtering (A), alignments trimmed with the automated heuristic column-based filtering of trimAl (B), and alignments requiring a full data matrix with no missing taxa per locus and with columns containing gaps or undetermined bases removed (C).

**Supplemental Figure 3.**
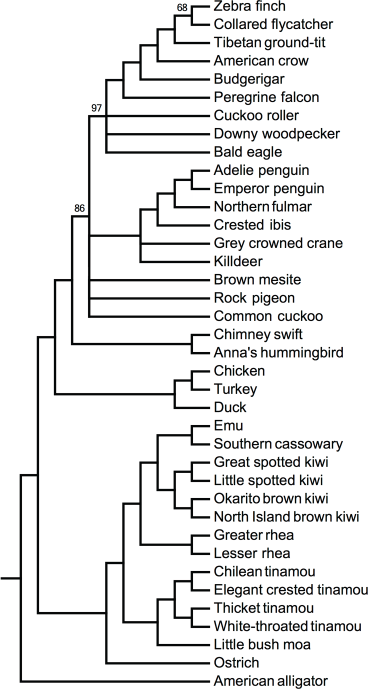
MP-EST species tree for 6,931 CNEEs. The recovered topology for palaeognaths is robust to the choice of an alternative outgroup (American alligator), and the inclusion of additional avian ingroup taxa. Branches with support < 50% are collapsed, and only bootstrap support values < 100% are drawn.

**Supplemental Figure 4.**
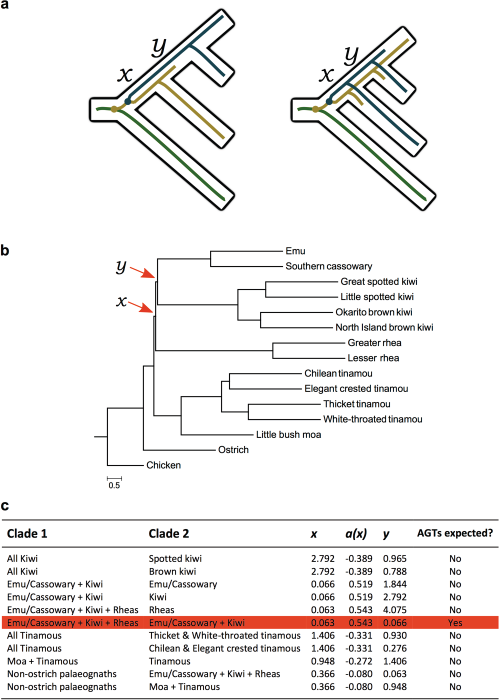
Evidence for incomplete lineage sorting. A) Incomplete lineage sorting (ILS) can produce anomalous gene trees (AGTs) across pairs of short internal branches, labelled x and y. Colored lines represent different alleles, and circles indicate mutational events. B) MP-EST species tree for the 20,850 locus total evidence data set, with internal branch lengths in coalescent units from MP-EST analysis of maximum likelihood RAxML gene trees. Terminal branch lengths are uninformative and are drawn as a constant value across all taxa. C) Branch lengths in coalescent units for all pairs of branches x and y across the species tree, with the boundary of the anomaly zone a(x) calculated following Equation 4 from Degnan and Rosenberg (Degnan and Rosenberg 2006). AGTs are expected when y < a(x).

**Supplemental Figure 5.**
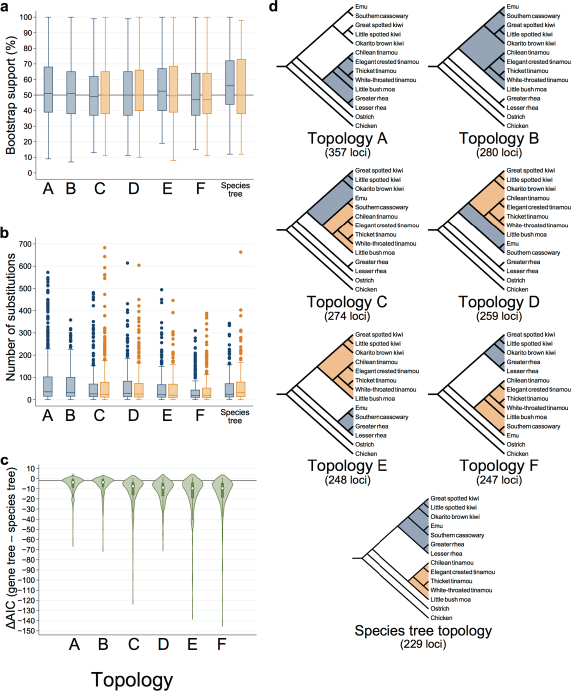
Gene trees that conflict with the MP-EST species tree have strong support. Boxplots of bootstrap support (A) and the number of substitutions under a parsimony criterion (B) for clades that conflict with the species tree topology. A reference line of 50% bootstrap support is drawn in (A). C) Violin plots of the difference in Akaike information criteria (AIC) for the recovered gene tree topology for each locus relative to the AIC when the sequence alignment is constrained to the species tree topology. A reference line is drawn at ΔAIC= -2, indicating substantial support in favor of the gene tree topology over that of the species tree for a given locus. D) Diagrams of the six gene tree topologies that occur at highest frequency (topologies A–F), and of the species tree topology. Clades considered in parts A and B are shaded and marked with star symbols.

**Supplemental Figure 6.**
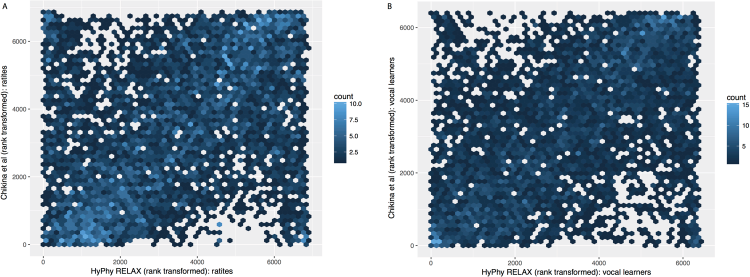
Consistent results from RELAX and branch tests. We compared the rank-transformed estimates of changes in constraint for both ratites (A) and vocal learners (B) between the amino acid relative rate test (Chikina et al. 2016) and the HyPhy RELAX test for changes in selection intensity. Despite operating on different data types and with radically different statistical approaches these two quantities are highly correlated: Spearman’s rho = 0.337 (ratites) and rho = 0.403 (vocal learners), in both cases with P < 2.2e-16. Note we expect some points in the upper left and lower right corners representing cases where RELAX detects changes in intensity of selection for a gene with dN/dS greater than 1, which will give reversed expectations in the relative rate test.

**Supplemental Figure 7.**
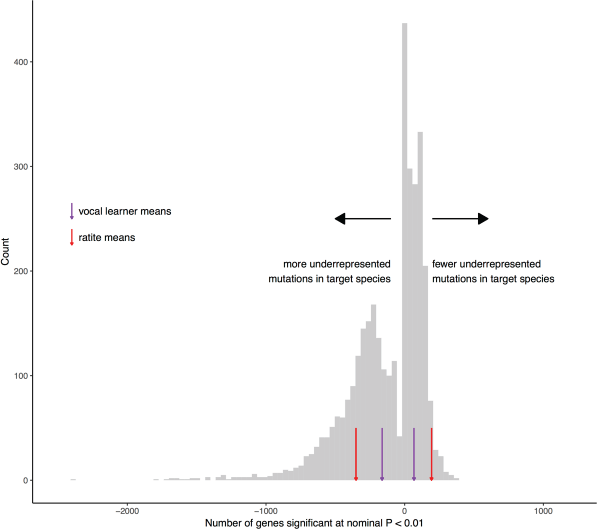
Neither ratites nor vocal learners are enriched for convergent, putatively function-altering substitutions in protein-coding genes. We sampled random trios of non-sister birds species (excluding trios that are entirely vocal learners or entirely ratites) and computed the number of proteins with significantly high or low DeltaBS scores for each trio compared to all other birds, as described in methods. Counts of proteins significant at a nominal P = 0.01 for both positive (amino acid sequence in trio is a better match to consensus than outside trio) and negative (amino acid sequence in trio is more diverged from the consensus than outside trio) DeltaBS scores are plotting for these random samples. Mean counts for ratite and vocal-learning trios (also excluding sister lineages) are plotted as vertical lines.

**Supplemental Figure 8.**
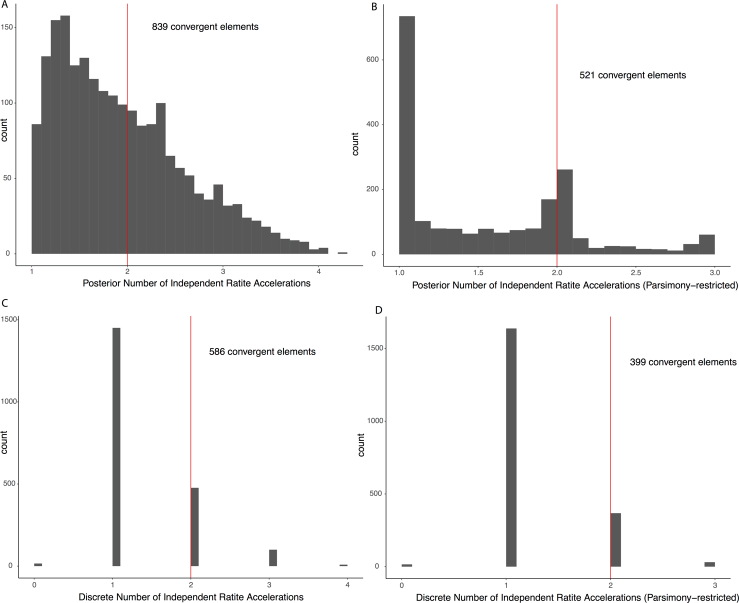
Ratite lineages experience frequent convergent acceleration in CNEEs. Histograms of estimates of independent (convergent) accelerations in ratites across all 2,053 RARs under different models. Red line shows cutoff at 2, above which we define a RAR as convergently accelerated. A) Posterior (continuous) estimate with no constraints on number of estimated losses per monophyletic clade. B) Posterior (continuous) estimate, but allowing a maximum of one acceleration event per monophyletic clade (e.g., up to one loss in ostrich, one loss in moa, and one loss in the rhea/emu/cassowary/kiwi clade). C) Discrete estimate produced by considering an element accelerated if the posterior estimate of accelerated on that branch is >= 0.95, conserved otherwise. D) Discrete estimate as in C, but with the additional parsimony constraint of B.

**Supplemental Figure 9.**
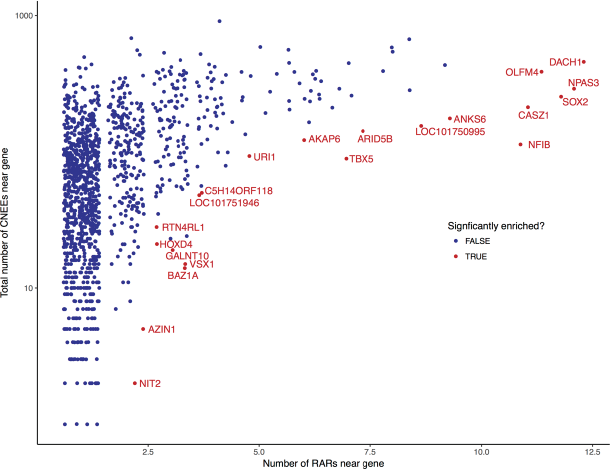
Genes with evidence for excess RARs. Each point represents the number of CNEEs associated with a gene and the number of RARs associated with that gene. Colored and labeled points are genes with excess RARs based on a permutation test (5% FDR). Only genes with at least one RAR are plotted.

**Supplemental Figure 10.**
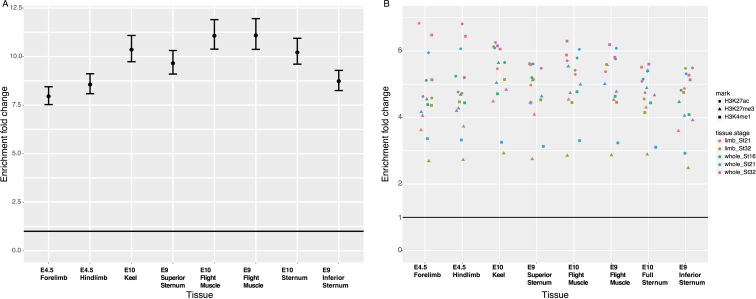
TSSs and ChIP-seq peaks are enriched under ATAC-seq peaks. A) TSSs are highly enriched under ATAC-seq peaks across all eight chicken tissues. Enrichment fold change was calculated against chicken genomic background using GAT after 10,000 samples. B) Previously published ChIP-seq peaks are also enriched under ATAC-seq peaks across all eight chicken tissues. Enrichment fold change was calculated against chicken genomic background using GAT after 10,000 samples.

**Supplemental Table 1:** Source information for Palaeognath genomes included in this study

| Code | Scientific name | Common name | Accession | Institution<sup>1</sup> | Collection locality | Sex | Reference |
| --- | --- | --- | --- | --- | --- | --- | --- |
| anoDid | *Anomalopteryx didiformis* | Little bush moa | TW95 | ROM | Pyramid Valley, Canterbury, New Zealand | unknown | Cloutier [27] |
| aptHaa | *Apteryx haastii* | Great spotted kiwi | R29945 | ROM | Hurunui, Canterbury, New Zealand | Male | This study |
| aptMan | *Apteryx mantelli* | North Island brown kiwi | multiple<sup>2</sup> | multiple<sup>2</sup> | see ref.<sup>2</sup> | Female | Le Duc [29] |
| aptOwe | *Apteryx owenii* | Little spotted kiwi | O 20599 | ROM | Te Mimiorakopa, Kapiti Island, New Zealand | Male | This study |
| aptRow | *Apteryx rowi* | Okarito brown kiwi | R 43461 | ROM | Okarito, Westland, New Zealand | Male | This study |
| casCas | *Casuarius casuarius* | Southern cassowary | B32419 | ANWC | Babinda, Queensland, Australia | Female | This study |
| cryCin | *Crypturellus cinnamomeus* | Thicket tinamou | 337707 | MCZ | Cafetal stream, Guanacaste, Costa Rica | Male | This study |
| droNov | *Dromaius novaehollandiae* | Emu | Cryogenic 6597 | MCZ | Songline Emu Farm, Gill, Massachusetts<sup>3</sup> | Male | This study |
| eudEle | *Eudromia elegans* | Elegant crested tinamou | 44215 | ROM | Toronto Zoo, Ontario, Canada | Male | This study |
| notPer | *Nothoprocta perdicaria* | Chilean tinamou | Ctin4 | ROM | Chilliwack, British Columbia, Canada<sup>3</sup> | Male | This study |
| tinGut | *Tinamus guttatus* | White-throated tinamou | B-42614 | LSU | Loreto Department, Peru | Female | Jarvis [28] |
| rheAme | *Rhea americana* | Greater rhea | MKP 2758 | ROM | Viedma, Rio Negro, Argentina | Male | This study |
| rhePen | *Rhea pennata* | Lesser rhea | 206185 (NZPBD100-12) | SNZ | Sea World, Orlando, Florida<sup>4</sup> | Male | This study |
| strCam | *Struthio camelus* | Ostrich | 202443 | SDZ | Botswana, Africa | Female | Jarvis [28] |

<sup>1</sup>ANWC: Australian National Wildlife Collection
MCZ: Harvard University Museum of Comparative Zoology
ROM: Royal Ontario Museum
SNZ: Smithsonian National Zoo
SDZ: San Diego Zoo

**Supplemental Table 2:** Sequencing and assembly information for new genomes

| Scientific name | Total Frag. Reads | Total MP Reads | Scaffold N50 | Contig N50 | BUSCO score<sup>1</sup> |
| --- | --- | --- | --- | --- | --- |
| *Apteryx haastii* | 294,924,492 | 277,444,152 | 1360 | 105.6 | C:97.1, F:2.0, M:0.9 |
| *Apteryx owenii* | 447,749,042 | 451,574,758 | 1617 | 128.4 | C:97.6, F:1.6, M:0.8 |
| *Apteryx rowi* | 357,835,928 | 436,303,038 | 1665 | 120.3 | C:98.1, F:1.3, M:0.6 |
| *Casuarius casuarius* | 419,359,436 | 459,568,992 | 3700 | 133.1 | C:97.8, F:1.3, M:0.9 |
| *Crypturellus cinnamomeus* | 269,962,512 | 322,211,994 | 2425 | 50.3 | C:97.2, F:1.4, M:1.4 |
| *Dromaius novaehollandiae* | 501,233,652 | 397,997,402 | 3322 | 138.8 | C:97.7, F:1.4, M:0.9 |
| *Eudromia elegans* | 342,759,376 | 329,360,988 | 3283 | 98.8 | C: 98.1, F: 1.1, M: 0.8 |
| *Nothoprocta perdicaria* | 429,304,722 | 336,279,414 | 3348 | 75.8 | C: 97.4, F: 1.8, M: 0.8 |
| *Rhea americana* | 390,468,074 | 375,784,406 | 4082 | 68.7 | C: 97.6, F: 1.2, M: 1.2 |
| *Rhea pennata* | 312,785,672 | 331,128,486 | 3846 | 55.9 | C: 96.3, F: 1.5, M: 2.2 |

<sup>1</sup>BUSCO scores are percentage of complete (C), fragmented (F), and missing (M) vertebrate BUSCO models in each genome.

**Supplemental Table 3:** Predicted genes in each genome

| Scientific name | # Gene models | % ortho (OMA) | % chicken blastp<sup>1</sup> | % eggNOG hit<sup>2</sup> | BUSCO score<sup>3</sup> |
| --- | --- | --- | --- | --- | --- |
| *Apteryx haastii* | 19,973 | 72.4% | 98.80% | 86.6% | C: 94.5, F: 4.3, M: 1.2 |
| *Apteryx owenii* | 20,969 | 70.9% | 99.07% | 86.1% | C: 94.6, F: 3.9, M: 1.5 |
| *Apteryx rowi* | 20,710 | 71.2% | 99.00% | 85.7% | C: 95.1, F: 3.6, M: 1.3 |
| *Casuarius casuarius* | 21,342 | 72.4% | 99.02% | 87.1% | C: 93.6, F: 5.2, M: 1.2 |
| *Crypturellus cinnamomeus* | 18,883 | 83.7% | 98.93% | 95.7% | C: 95.3, F: 3.8, M: 0.9 |
| *Dromaius novaehollandiae* | 18,858 | 77.0% | 98.90% | 89.2% | C: 96.1, F: 3.1, M: 0.8 |
| *Eudromia elegans* | 17,021 | 84.9% | 98.77% | 95.6% | C: 95.4, F: 3.7, M: 0.9 |
| *Nothoprocta perdicaria* | 18,151 | 78.5% | 98.91% | 88.8% | C: 96.1, F: 2.9, M: 1.0 |
| *Rhea americana* | 17,916 | 81.6% | 98.61% | 94.3% | C: 94.0, F: 4.9, M: 1.1 |
| *Rhea pennata* | 19,203 | 80.5% | 98.88% | 94.1% | C: 93.8, F: 5.4, M: 0.8 |

<sup>1</sup>Percent of chicken proteins with a blast hit to target species predicted proteome.
<sup>2</sup>Percent of predicted proteins in target species with a hit to a vertebrate eggNOG model.
<sup>3</sup>BUSCO scores are percentage of complete (C), fragmented (F), and missing (M) vertebrate BUSCO models in each proteome.

**Supplemental Table 4:** Phylogenomic data sets used for MP-EST coalescent and ExaML concatenation species tree inference

| Marker | Data set | Num. loci | Aligned sequence length (bp) |
| --- | --- | --- | --- |
| **CNEEs** | MAFFT | 12,676 | 4,794,620 |
|  | MAFFT trimAl | 12,676 | 4,667,306 |
|  | MAFFT no missing data | 11,125 | 3,851,232 |
|  | MAFFT allspecies (American alligator outgroup) | 6,931 | 2,937,637 |
| **Introns** | MAFFT | 5,016 | 27,890,802 |
|  | MAFFT trimAl | 5,016 | 23,346,653 |
|  | MAFFT no missing data | 2,117 | 2,333,861 |
| **UCEs** | MAFFT | 3,158 | 8,498,759 |
|  | MAFFT trimAl | 3,158 | 7,361,135 |
|  | MAFFT no missing data | 1,837 | 2,409,470 |
| **TENT** | MAFFT | 20,850 | 41,184,181 |
|  | MAFFT trimAl | 20,850 | 35,375,094 |
|  | MAFFT no missing data | 15,079 | 8,594,563 |

**Supplemental Table 5:** Model fits with number of convergent substitutions as response variable

| Data Filtering | # Alignments | Model Terms | Estimate<sup>1</sup> | P-value |
| --- | --- | --- | --- | --- |
| **Default**<br>(1:1 orthologs only,<br>< 3 missing taxa, species tree) | 6,337 | Divergent Substitutions | 0.7885 | < 2e-16 |
|  |  | Evolutionary Distance | -766.97 | 0.00146 |
|  |  | Divergent:Distance interaction | -1.01 | < 2e-16 |
|  |  | **Is Ratite?** | **46.39** | **0.60375** |
|  |  | *Overall* | 0.9767 | < 2e-16 |
| **Default**<br>(1:1 orthologs only,<br>< 3 missing taxa, species tree) | 6,337 | Divergent Substitutions | 0.1113 | 6.43e-11 |
|  |  | Log(Evolutionary Distance) | -126.8 | 0.0284 |
|  |  | Divergent:Distance interaction | -0.3035 | < 2e-16 |
|  |  | **Is Ratite?** | **68.82** | **0.4818** |
|  |  | *Overall* | 0.9778 | < 2e-16 |
| **Default**<br>(1:1 orthologs only,<br>< 3 missing taxa, species tree) | 6,337 | Divergent Substitutions | 1.4676 | < 2e-16 |
|  |  | Exp(Evolutionary Distance) | -580.28 | 0.00172 |
|  |  | Divergent:Distance interaction | -0.724 | < 2e-16 |
|  |  | **Is Ratite?** | **52.54** | **0.55808** |
|  |  | *Overall* | 0.9757 | < 2e-16 |
| **Gene Tree**<br>(1:1 orthologs only,<br>< 3 missing taxa, gene tree) | 6,342 | Divergent Substitutions | 0.6981 | < 2e-16 |
|  |  | Evolutionary Distance | -620.6 | 3.9e-07 |
|  |  | Divergent:Distance interaction | -0.8271 | < 2e-16 |
|  |  | **Is Ratite?** | **-5.018** | **0.9054** |
|  |  | *Overall* | 0.9809 | < 2e-16 |
| **Gene Tree**<br>(1:1 orthologs only,<br>< 3 missing taxa, gene tree) | 6,342 | Divergent Substitutions | 0.1171 | < 2e-16 |
|  |  | Log(Evolutionary Distance) | -752.2 | 0.00266 |
|  |  | Divergent:Distance interaction | -0.2672 | < 2e-16 |
|  |  | **Is Ratite?** | **31.23** | **0.47443** |
|  |  | *Overall* | 0.982 | < 2e-16 |
| **Gene Tree**<br>(1:1 orthologs only,<br>< 3 missing taxa, gene tree) | 6,342 | Divergent Substitutions | 1.241 | < 2e-16 |
|  |  | Exp(Evolutionary Distance) | -501.7 | 1.79e-07 |
|  |  | Divergent:Distance interaction | -0.5837 | < 2e-16 |
|  |  | **Is Ratite?** | **-5.576** | **0.896** |
|  |  | *Overall* | 0.9798 | < 2e-16 |
| **Strict**<br>(1:1 orthologs only,<br>no missing taxa, gene tree) | 3,946 | Divergent Substitutions | 0.7204 | < 2e-16 |
|  |  | Evolutionary Distance | -380.84 | 1.42e-06 |
|  |  | Divergent:Distance interaction | -0.852 | < 2e-16 |
|  |  | **Is Ratite?** | **-3.15** | **0.9080** |
|  |  | *Overall* | 0.9804 | < 2e-16 |
| **Strict**<br>(1:1 orthologs only,<br>no missing taxa, gene tree) | 3,946 | Divergent Substitutions | 0.1232 | < 2e-16 |
|  |  | Log(Evolutionary Distance) | -45.81 | 0.00544 |
|  |  | Divergent:Distance interaction | -0.2741 | < 2e-16 |
|  |  | **Is Ratite?** | **19.7** | **0.49144** |
|  |  | *Overall* | 0.9811 | < 2e-16 |
| **Strict**<br>(1:1 orthologs only,<br>no missing taxa, gene tree) | 3,946 | Divergent Substitutions | 1.281 | < 2e-16 |
|  |  | Exp(Evolutionary Distance) | -307.56 | 6.28e-07 |
|  |  | Divergent:Distance interaction | -0.6025 | < 2e-16 |
|  |  | **Is Ratite?** | **-3.454** | **0.9** |
|  |  | *Overall* | 0.9794 | < 2e-16 |

<sup>1</sup>Estimate is model coefficient for model terms, and the adjusted R-squared for the entire model for the overall term.

